# Information coding strategies in biological reservoir computing for multiclass time-series classification

**DOI:** 10.64898/2026.10.22.755221

**Authors:** Yusei Nishi, Hideaki Yamamoto, Yuichi Katori, Shigeo Sato

## Abstract

Physical reservoir computing (RC) using biological neural networks (BNNs) offers a framework for probing how high-dimensional dynamics arising from multicellular systems support complex information processing in the brain. However, how electrical inputs and extracellular readouts should be designed to suit different task demands remains unclear. Here, using primary rat cortical neurons cultured on a high-density microelectrode array, we investigated how input encoding and spike detection methods influence neural representations and the resulting RC performance in BNNs. Comparison of pulse amplitude modulation and latency coding revealed that the latter exhibited superior performance in demanding multiclass conditions, including a 10% improvement in a 10-class spoken digit classification task. Spike sorting further improved classification performance by enhancing class separability, while largely preserving the low-dimensional structure of population activity. These findings highlight the importance of designing information coding strategies according to task difficulty.

## 1 Introduction

Biological brains exploit a complex network of neurons and their high-dimensional population dynamics as a physical substrate to process information and enable functions such as sensory perception and behavioural control [1–3]. This perspective has motivated the use of recurrent neural networks (RNNs) as models for such temporal information processing in the brain [4]. Previous studies have shown that trained RNNs can perform temporal cognitive tasks associated with the prefrontal cortex, such as decisionmaking and working memory [5–7]. As an engineering application of RNNs in vivo, the brain–machine interface exploits the neural population dynamics generated by recurrent neural dynamics in the brain and translates them into control signals for external devices [8, 9].

Although powerful, RNNs suffer from the heavy computational demand required to optimize the recurrent weights in response to temporal signals. Reservoir computing (RC) has since been proposed as a paradigm that uses RNNs with fixed internal weights to map inputs into high-dimensional state trajectories from which task-relevant information is detected by a simple, memory-less linear decoder [10–13]. Notably, RC is, at least in part, consistent with in vivo observations that recurrently connected biological neural networks (BNNs) encode behaviourally relevant variables through population dynamics in the prefrontal cortex and secondary motor cortex [14, 15]. Moreover, BNN-based RCs have attracted attention as a framework for bottom-up investigations of the relationship between neural dynamics and information processing [16–20]. Such in vitro BNNs have emerged as controllable physical platforms for tracing how cellular-scale circuit organization gives rise to population dynamics that can be harnessed for computation and have been applied to various computational tasks, including time-series prediction and generation [19, 20], spatial pattern classification [21, 22], and closed-loop learning [17, 20, 23]. Among these tasks, spoken digit classification is particularly suitable for evaluating reservoir-like temporal processing because speech signals contain rich temporal structures and require nonlinear transformation and short-term memory [18, 19].

With respect to physical RC using in vitro BNNs, neurons are typically interfaced either electrically using a microelectrode array (MEA) [24–26] or optically using techniques such as calcium imaging and optogenetics [18]. Among these approaches, MEAs offer high temporal resolution and direct, noninvasive access to electrophysiological activity, making them particularly suitable for investigating neuronal network dynamics for RC. To implement RC tasks in BNNs interfaced with MEAs, input signals must be encoded into spatiotemporal patterns of electrical stimulation, while recorded extracellular potentials must be decoded to generate task-relevant outputs. Various input encoding methods, such as pulse amplitude modulation (PAM), pulse rate modulation, and timing-based coding [19, 20, 25, 27], together with spike detection methods, including threshold-based detection and spike sorting [28, 29], have been explored to deliver inputs to BNNs and read out their responses. However, a systematic investigation of how these components influence neural representations and RC performance remains lacking, and how they should be designed to suit different task demands is poorly understood.

Here, we evaluated how information coding strategies influence neural representations and computational performance in BNN-based RC. To this end, we implemented a spoken digit classification task in primary rat cortical networks grown on a high-density microelectrode array (HD-MEA). We compared two input encoding schemes, namely, PAM [20, 30] and latency coding (LC) [19], and further compared threshold-based detection [28] and spike sorting [29] as methods for detecting spikes. Our results show that input encoding schemes determine how information is represented in neural activity. Moreover, spike sorting enhanced RC performance without substantially altering the global structure of population activity. However, these benefits were limited in relatively less demanding classification tasks, suggesting that information coding strategies in BNN-based RC systems should be designed according to task complexity.

## 2 Methods

### 2.1 Microfluidic device

Microfluidic devices for patterning neurons were constructed according to a previously reported protocol [31, 32]. In brief, SU-8 photoresists (Kayaku Advanced Materials) were patterned on a silicon wafer to prepare a master mould. Polydimethylsiloxane (PDMS; Sylgard 184 mixed at a 10:1 ratio) was subsequently cast from the edge of the mould and cured at 70 °C for 2 h. After curing, the PDMS was detached from the mould and cut with a clean razor blade. The device was then cleaned by sonication in ethanol for 15 min, washed three times with deionized water, and sterilized by UV exposure for 30 min. The device was then attached to the recording area of the HD-MEA chip.

### 2.2 Cell culture

Primary rat cortical neurons were obtained from embryonic Day 18 rat embryos following a previously published protocol [20, 30]. Before cell plating, the electrode surface was treated with 1% terg-a-zyme (Alconox 1304-1) for hydrophilization and incubated at room temperature for 2 h, followed by three rinses with deionized water. Next, the chip was submerged in a beaker filled with 70% ethanol for sterilization and left for 30 min. The chip was then rinsed three times with deionized water and dried. The electrode surface was subsequently covered with a 0.07% PEI/borate buffer solution and incubated for 1 h. The surface was then rinsed with deionized water and dried. Finally, a 50–100 µg/mL laminin/DPBS solution was applied using the same procedure as for the PEI coating and the electrode surface was rinsed three times with deionized water and dried.

Following the coating, a microfluidic device was attached to the electrode and left overnight to allow firm adhesion. The chip well was then filled with neuronal plating medium (minimum essential medium (MEM) (Gibco 11095-080) + 5% foetal bovine serum + 0.6% D-glucose (Sigma G-8769)), and the air bubbles trapped in the microfluidic device were removed using a vacuum chamber. Afterwards, the medium was completely replaced with fresh plating medium and incubated in a CO_2_ incubator (37 °C, 5% CO_2_) for at least 3 h. Cortical neurons were plated at a density of 700 cells/mm^2^ and incubated for 1 h. Finally, the medium was completely replaced with Neurobasal medium (Neurobasal medium (Gibco 21103-049) + 2% B-27 (Gibco 17504-044) + 1% GlutaMAX-I (Gibco 35050-061)), and half the medium was changed twice per week.

Neuronal growth on HD-MEA was assessed by the fluorescent chemical probe NeuO [33] as described previously ([20]). Briefly, cells were incubated with 2 µM (STEMCELL Technologies 01801) in Neurobasal medium at 37 °C for 30 min, and the medium was then replaced with NeuO-free medium. Images were captured with an sCMOS camera (Zyla 4.2P; Andor) mounted on a stereomicroscope (SMZ18; Nikon).

### 2.3 Spoken digit classification task using HD-MEA

Neural recordings were performed using the MaxOne HD-MEA system (MaxWell Biosystems). The MaxOne+ chip contains 26,400 electrodes arranged at a pitch of 17.5 µm within a sensing area of 3.85 ×2.10 mm^2^. Extracellular potentials were recorded at a sampling rate of 20 kHz and processed offline using the SpikeInterface package [34].

Spoken digit classification was used as the benchmark task. The TI-46 spoken dataset consisting of spoken digits from “zero” to “nine” spoken by eight speakers, including four males and four females, was used. For each speaker and digit class, three trials were used for training and one trial was used for testing. In analysis involving fewer than 10 classes, classification accuracy was averaged over all possible combinations of digit classes.

Input signals were converted into electrical stimulation sequences using two encoding methods, namely, PAM [20, 30] and LC [19]. For both methods, one stimulation electrode was selected from an active region in each of the 30 wells from the 128 wells. Electrical stimulation was delivered as biphasic pulses, and the pulse width was set to 200 µs per phase. Each stimulation sequence was presented with an interval of 15 s For PAM, a 3-dimensional cochleagram was generated using Lyon’s passive ear model, and each dimension was assigned to a group of 10 spatially adjacent electrodes selected from the 30 stimulation electrodes. For each trial, the cochleagram amplitudes were scaled to the range [0, 1]. The resulting normalized amplitude 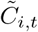 in dimension *i* at time *t* was linearly mapped onto the amplitude *A*_*i,t*_ of a biphasic stimulation pulse according to

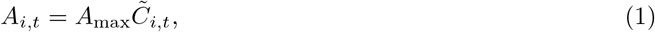

where *A*_max_ = 900 mV. The stimuli were delivered at intervals of 50 ms. Although increasing the dimensionality of the cochleagram used for PAM may improve performance, the PAM implementation was limited to three dimensions under the present MEA stimulation setup. Therefore, the configuration used here was selected to maximize the performance for PAM within the constraints of the current system.

For LC, a 30-dimensional cochleagram was generated using the same procedure as for PAM, and each dimension was assigned to a unique stimulation electrode. The stimulation amplitude was fixed at 650 mV, which evoked stable responses [20]. Each cochleagram frame was assigned to a 100 ms stimulation window, and the delay Δ_*i,t*_ after the beginning of each window was determined by

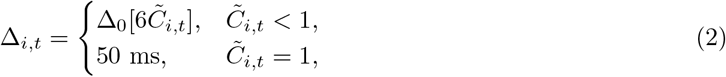

where Δ_0_ = 10 ms and [·] denotes the floor function. This mapping yielded stimulation delays ranging from 0 to 50 ms in increments of 10 ms. Electrical stimulation artefacts were removed from the extracellular traces by excluding data within a 3 ms window around each stimulation timing and reconstructing the removed segments by third-order spline interpolation. The signal was bandpass filtered between 300 and 3000 Hz, and a common median reference [35] was applied using a local reference with a spatial radius of 400 µm.

Spike events were then detected from the preprocessed traces using either threshold-based detection or spike sorting. For threshold-based detection, the root mean square (RMS) of the preprocessed extracellular traces was computed at each electrode, and spikes were subsequently detected when the magnitude of a negative peak exceeded seven times the RMS value. Spike sorting was performed using Kilosort4 [29], and only clusters labelled as “good” by the Kilosort4 criteria were used in the analysis.

Spike trains obtained by these detection methods were binned with a temporal resolution of 10 ms to construct a matrix **X** ∈ℝ^*T ×d*^, where *T* and *d* denote the time points and the number of units, respectively. Here, *d* corresponds to the number of electrode channels for threshold-based detection and the number of clusters for spike sorting. The matrix was then flattened into a one-dimensional activity vector. Because this representation preserves the absolute temporal positions of neural activity, it is sensitive to temporal misalignment across input trials. Specifically, even within the same digit class, temporal variations in the input cochleagrams may shift the resulting neural activity across time bins, potentially degrading classification performance. To reduce this effect, a large number of training trials were used. These activity vectors were used for classification, where logistic regression implemented in scikit-learn was used to train the readout weights and perform classification.

### 2.4 Principal component analysis

The low-dimensional structure of neural population activity was characterized by principal component analysis (PCA) [20]. For this analysis, **X** was centred by subtracting the temporal mean of each unit to obtain the centred firing rate matrix 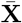. The covariance matrix **Σ** was then computed as follows:

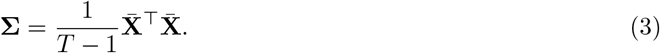

The factor loadings and explained variances were derived through eigendecomposition:

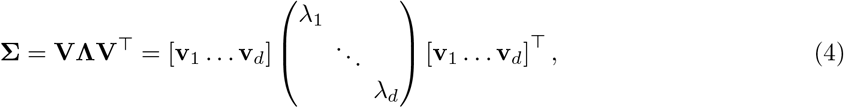

where **V** and **Λ** represent the eigenvector matrix and the diagonal matrix of eigenvalues *λ*_*i*_, respectively. The eigenvectors **v**_*i*_ correspond to the factor loadings of the *i*-th PC and each normalized eigenvalue *p*_*i*_ = *λ*_*i*_/ (Σ_*j*_ *λ*_*j*_) represents the variance explained by the corresponding PC. The effective dimensionality of neural population activity was then quantified using the effective rank as follows: erank(**X**) = exp (*H*(*p*_1_, …, *p*_*d*_)), where *H* is the Shannon entropy [36].

To evaluate whether population-level dynamics were preserved across spike detection methods, canonical correlation analysis (CCA) was performed to determine the correlation between the activity obtained by threshold-based detection and spike sorting [37]. Specifically, the CCA yields linear projections of two projected activity trajectories such that the correlation between the corresponding projected trajectories is maximized. The resulting canonical correlations were used to quantify how well the activity trajectories obtained by the two detection methods could be aligned in a shared dimensional space.

For each recording, the binned spike trains were smoothed using a double exponential kernel with rise and decay time constants of 50 ms and 200 ms, respectively. The smoothed firing rate matrices obtained by threshold-based detection and spike sorting are denoted as follows: 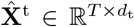 and 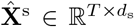, where *d*_t_ and *d*_s_ denote the number of electrode channels and sorted clusters, respectively. To prepare the activity trajectories for the CCA, PCA was performed separately on each smoothed firing rate matrix. The number of retained PCs was set to *d*_s_(≤ *d*_t_) so that the two PCA-projected trajectories had the same dimensionality. The resulting latent trajectories are denoted as 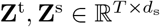. For the *k*-th aligned canonical dimension (neural mode), the trajectories obtained by threshold-based detection and spike sorting were linearly projected as follows:

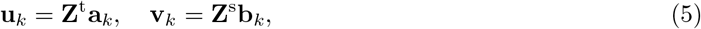

where **a**_*k*_ and **b**_*k*_ were chosen to maximize the correlation between **u**_*k*_ and **v**_*k*_. Finally, the canonical correlation coefficient was defined as follows:

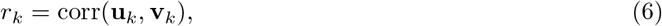

and used as a measure to quantify how well the two latent trajectories were aligned along the *k*-th aligned canonical dimension. As a control, one latent trajectory was shifted by 5 min relative to the other before the CCA was applied. This time-shifted control disrupts the time-point-wise correspondence between activity obtained by threshold-based detection and spike sorting while preserving the temporal structure within each trajectory.

### 2.5 Similarity analysis

For each trial, an activity vector was constructed by vectorizing the firing rate matrix **X**. The similarity between two activity vectors **a** and **b** was quantified using cosine similarity:

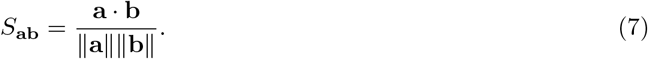

For each digit class, the class centroid was defined as the mean of the corresponding activity vectors. Interclass similarity was defined as the mean cosine similarity between the centroids of different classes, whereas intraclass similarity was calculated as the mean cosine similarity between each activity vector and the centroid of its corresponding class.

## 3 Results

### 3.1 Spontaneous activity of BNNs

We used a 10-class spoken digit classification task as a benchmark for evaluating the computational capacity of BNNs (Fig. 1a). Rat cortical neurons were cultured on an HD-MEA. All the experiments were conducted at 15–30 days in vitro (DIV). A microfluidic device was used to control the spatial arrangement of the neurons and their network connectivity on the HD-MEA surface to prepare a BNN that can generate rich population dynamics (Fig. 1b) [20, 31, 38]. The device consisted of 128 square wells with a side of 100 µm, each of which contained an average of 15 neurons [20]. The wells were arranged in an 8-by-16 lattice with an interwell spacing of 100 µm, and adjacent wells were interconnected by microtunnels of approximately 5 µm in both width and height. Spoken digit inputs were converted into electrical stimulation sequences using either LC or PAM and delivered to the BNN. The evoked neural activity was then detected using either threshold-based detection or spike sorting, and the resulting reservoir states were represented as activity vectors for digit classification. Detailed descriptions are provided in the Methods.

**Fig. 1.**
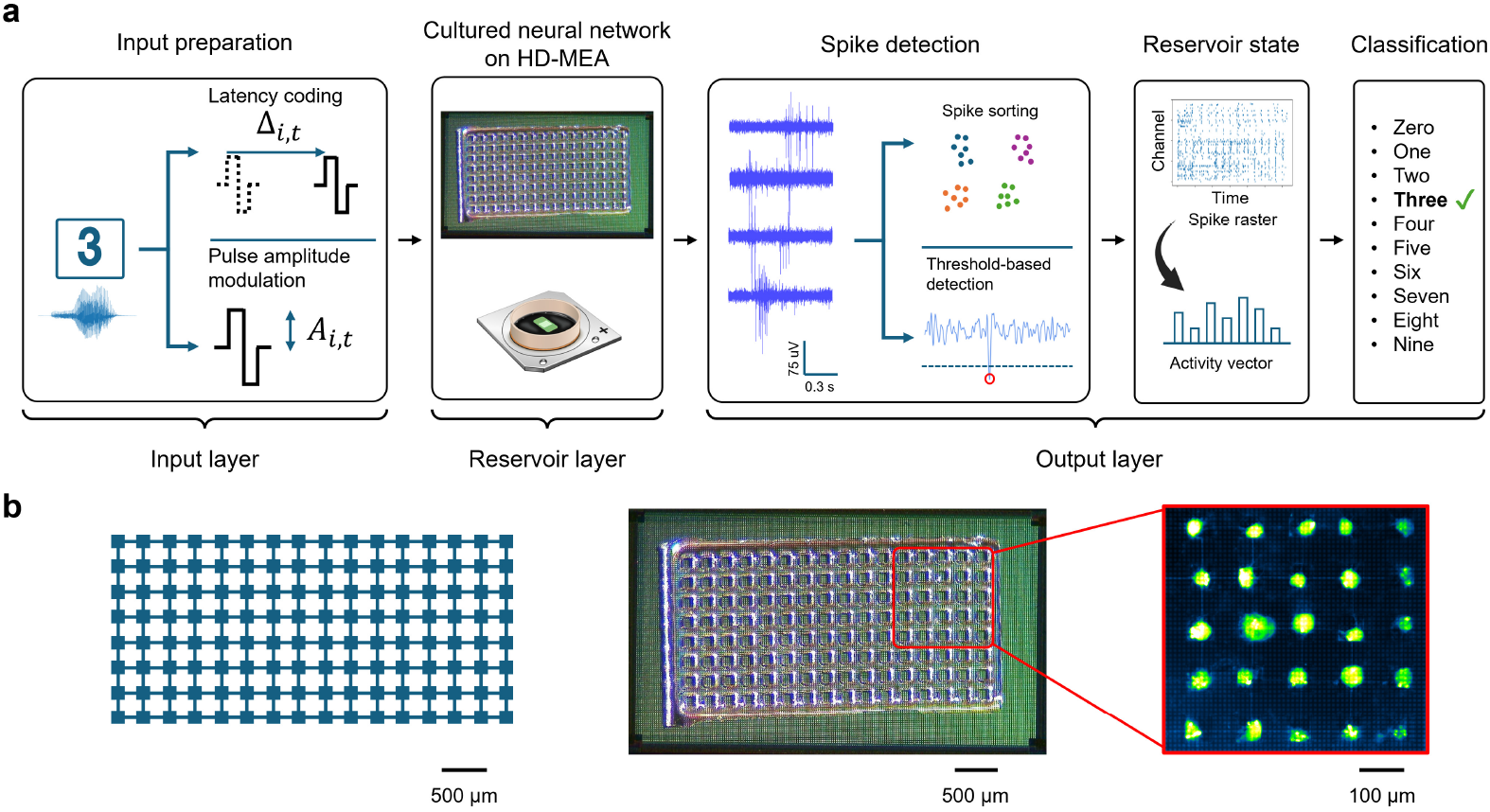
Overview of the experimental setup. (a) Spoken digit classification task. Input signals were coded into electrical stimulation sequences using either latency coding or pulse amplitude modulation and delivered to a cultured neuronal network on an HD-MEA. Spike events were detected from the evoked extracellular voltage traces using either spike sorting or threshold-based detection. The resulting spike trains were then binned and flattened to construct activity vectors. These activity vectors were used as inputs to logistic regression for classification. (b) Micropatterned cultured neuronal network constructed on an HD-MEA. Shown are the lattice network design (left), the recording area of the HD-MEA with the PDMS-microfluidic devices attached (middle), and primary cortical neurons cultured in the lattice pattern and stained with NeuO at DIV 30 (right).

We began by characterizing the BNNs that served as the reservoir layer. Because the reservoir state is defined by the spiking events detected from extracellular potentials, we compared two widely used detection methods, i.e., threshold-based detection and spike sorting, on the features of the recorded spontaneous activity. Consistent with previous studies [20, 31, 38], the neural activity obtained by threshold-based detection in the 128 lattice networks exhibited reduced global synchrony compared with that in homogeneous networks, resulting in a mixture of globally and locally synchronized activity patterns (Fig. 2a). This activity was characterized by a mean firing rate of 0.74 Hz and a mean STTC of 0.08 (*n* = 16 recordings from 13 networks).

**Fig. 2.**
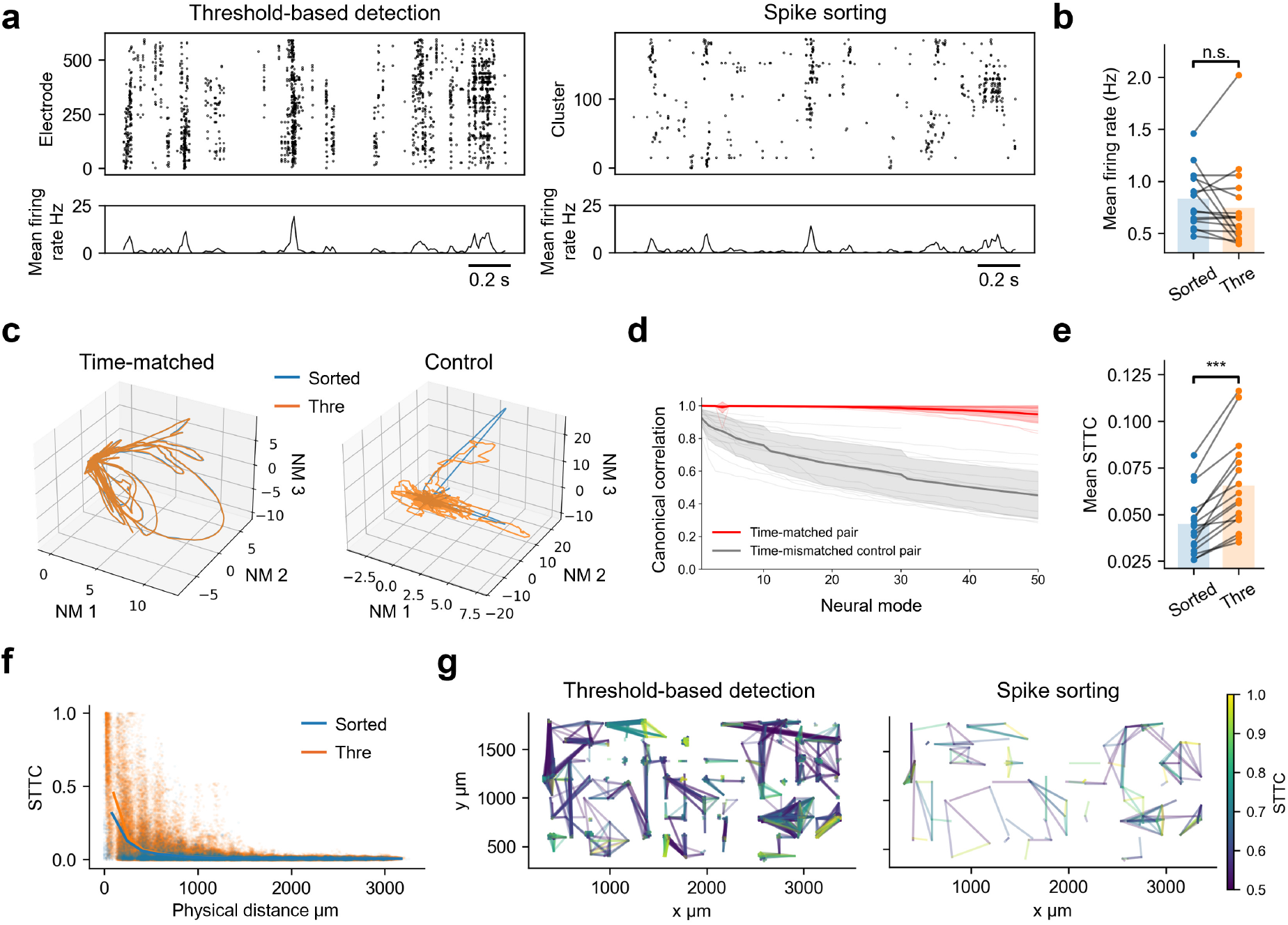
Effects of detection methods on analysis during spontaneous activity. (a) Representative spike rasters and firing-rate traces obtained by threshold-based detection (Thre; left) and spike sorting (Sorted; right). (b) Comparison of mean firing rates between threshold-based detection and spike sorting. Bars represent mean values in mean firing rate for each detection method. n.s., not significant (two-sided paired *t* test). (c) State trajectories of spontaneous activity projected onto the top three neural modes obtained by PCA followed by CCA. Threshold-based detection and spike sorting are shown in orange and blue, respectively. (d) Canonical correlations across neural modes. Time-matched pairs and time-mismatched control pairs are shown in red and grey, respectively. (e) Comparison of mean STTC between two detection methods. Bars represent mean values in STTC for each detection method. \*\*\**p <* 0.001 (two-sided paired *t* test). (f) Relationship between physical distance and STTC. Dots indicate individual electrode pairs, and lines indicate mean STTC. Data obtained using threshold-based detection and spike sorting are shown in orange and blue, respectively. (g) STTC networks constructed from electrode pairs with STTC *≥*0.5. Edges indicate electrode pairs exceeding the STTC threshold by threshold-based detection (left) and spike sorting (right). Sample size: *n* = 16 recordings from 13 networks for each detection method.

Analysis revealed that the two detection methods yielded spike rasters with comparable mean firing rates (Fig. 2a,b; *n* = 16 recordings from 13 networks; *p >* 0.05, two-sided paired *t* test). The latent dynamics, defined as low-dimensional trajectories of population activity, were compared after dimensionality reduction using PCA followed by dimensional alignment with CCA between the two detection methods. In the time-matched comparison, the aligned trajectories closely overlapped in the first three aligned dimensions, whereas the control pairs showed less overlap (Fig. 2c). The correlations were consistently higher for the time-matched pairs than for the control pairs, exceeding 0.9 for all of the first 50 aligned dimensions (Fig. 2d).

Next, we examined pairwise correlations using the spike time tiling coefficient (STTC) [39] with a time window of 10 ms. Pairwise correlations were significantly reduced by spike sorting, with a pronounced decrease in highly correlated pairs between spatially nearby electrodes (Fig. 2e, f). This reduction is consistent with the possibility that, in HD-MEA recordings, threshold-based detection samples the activity of a single neuron on multiple neighbouring electrodes, thereby inflating correlations among nearby channels. This is plausible because the electrodes are arranged at a 17.5 µm pitch, which is comparable to the size of a neuronal soma. Consequently, spike sorting led to sparser reconstructed functional networks (Fig. 2g), which may facilitate the identification of prominent functional networks. Collectively, these analyses suggest that spike sorting mainly involves integrating redundant detection of the same neuron across neighbouring electrodes, with little impact on population dynamics.

### 3.2 Effect of Encoding Methods

Input encoding methods can shape neural representations in the BNN reservoir and, hence, its task performance. To test this, we compared two approaches for converting input signals into electrical stimulation sequences: PAM [20, 30] and LC [19] (Fig. 3a). In PAM, the input signal modulated the amplitude of stimulation pulses delivered at a constant interval. In LC, the input signal modulated the timing of each stimulation pulse delivered at a constant amplitude (see Methods section for details). Under both conditions, evoked spike trains were obtained by spike sorting and used to construct activity vectors, which were then fed to a logistic regression for readout (Fig. 1a).

**Fig. 3.**
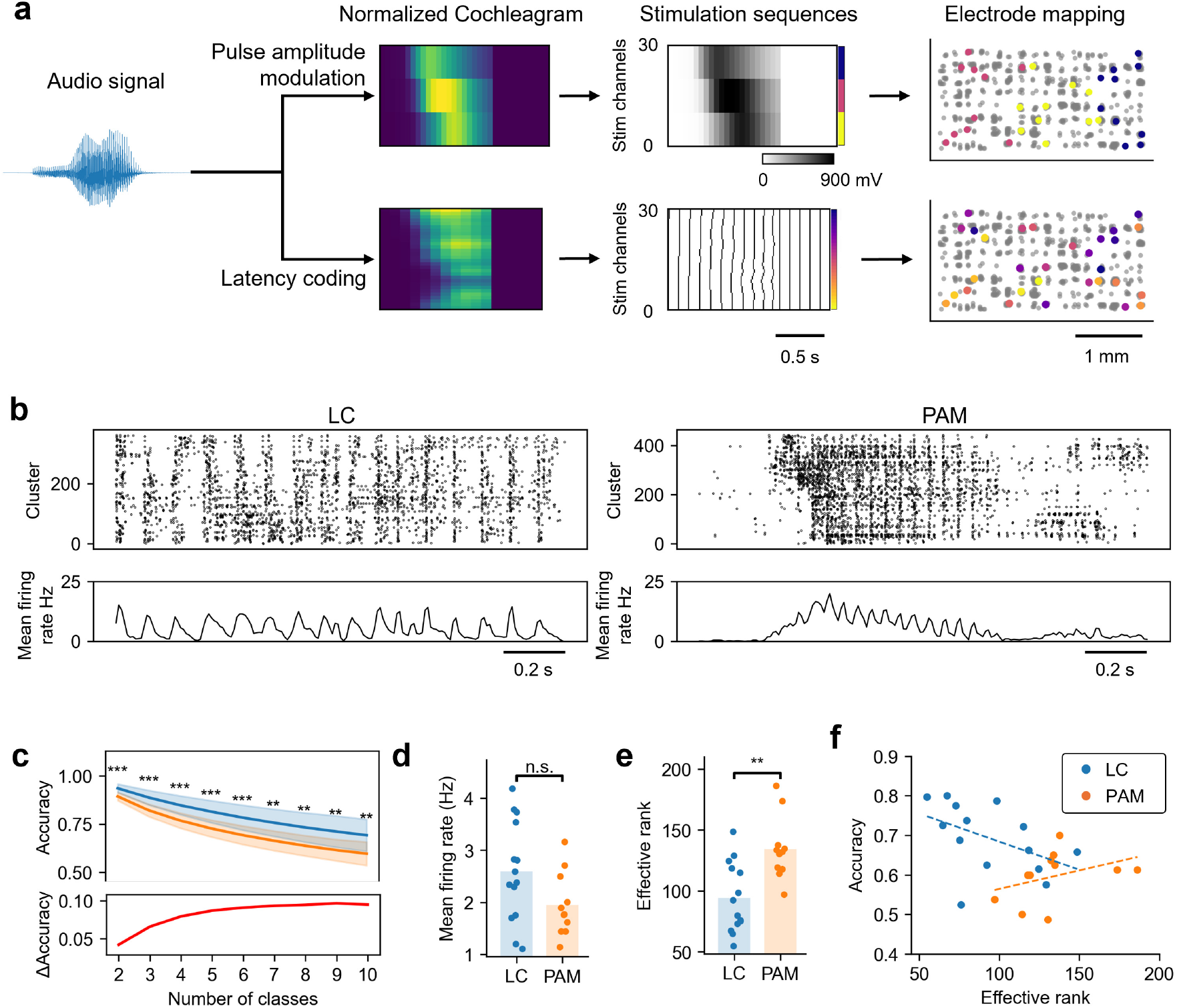
Effects of input encoding methods on reservoir computing. (a) Schematic overview of input signal encoding using LC and PAM. An audio signal was first converted into a cochleagram using Lyon’s cochlear filter. A cochleagram was then normalized to the range from 0 to 1. A normalized cochleagram was converted into electrical stimulation sequences using either LC or PAM, and the stimulation was delivered through the selected electrodes, as shown in the electrode maps on the right. (b) Representative spike rasters and firing-rate traces evoked by LC (left) and PAM (right). (c) Classification accuracy as a function of the number of input classes. Activity vectors obtained from each encoding method were used for classification by logistic regression. PAM and LC are shown in orange and blue, respectively. Lines indicate mean accuracy, and shaded areas indicate SD (top). The difference in mean accuracy between LC and PAM is presented in the bottom panel. \*\**p <* 0.01 and \*\*\**p <* 0.001 (two-sided Welch’s *t* test). (d) Comparison of mean firing rates between two encoding methods. Bars indicate the mean firing rates for each encoding method. n.s., not significant (two-sided Welch’s *t* test). (e) Comparison of effective rank between two encoding methods. Bars indicate the mean effective rank for each encoding method. \*\**p <* 0.01 (two-sided Welch’s *t* test). (f) Relationship between effective rank and classification accuracy. Orange and blue indicate PAM and LC, respectively. Dashed lines indicate linear fits for each encoding method. Sample size: *n* = 11 recordings from 10 networks (PAM), *n* = 14 recordings from 12 networks (LC).

Experiments revealed that the two methods generated distinct stimulation sequences and consequently induced different neural response patterns (Fig. 3b). In PAM, the input signal created a large transient increase in the population firing rate, whereas in LC, the population firing rate exhibited a smaller periodic oscillation at 10 Hz corresponding to the code interval of 100 ms. When the obtained spiking activity was decoded by a logistic regression, LC achieved significantly higher classification accuracy than PAM across all numbers of classes (Fig. 3c; PAM, *n* = 11 recordings; LC, *n* = 14 recordings). In the 10-class classification task, LC achieved an accuracy of 69.2 ±8.3% compared with 59.2 ±6.0% for PAM. However, the difference in classification accuracy was relatively smaller for tasks involving fewer classes and became more pronounced as the number of classes increased. The mean firing rate during stimulation was not significantly different between the two encoding methods (Fig. 3d; *p >* 0.05, two-sided Welch’s *t* test). This suggests that the differences in performance cannot be explained simply by variations in the overall activity level.

Interestingly, while high-dimensional dynamics are typically considered the basis of the computational performance of reservoir computing [40], the effective rank of the neural responses to the PAM-based stimulation was significantly higher than that obtained by the LC-based stimulation (Fig. 3e; *p <* 0.01, two-sided Welch’s *t* test), with both conditions exhibiting an effective rank sufficient for the present 10-class classification task. We therefore evaluated the relationship between the effective rank and classification performance for each encoding method (Fig. 3f). With respect to PAM, the classification accuracy showed a weak positive relationship with the effective rank (*r* = 0.37). In contrast, LC exhibited the opposite trend, with higher classification accuracy observed at lower effective ranks (*r* = −0.47). These results may reflect the rate-coding-like property of PAM and the temporal-coding-like property of LC. Thus, how the dimensionality of the reservoir state contributes to the classification performance depends on the input encoding method.

### 3.3 Effect of Spike Detection Methods

To evaluate how spike detection methods affect neural responses and classification performance in BNN-based reservoir computing, we compared threshold-based detection and spike sorting. For each method, spike trains were converted into activity vectors and used for logistic regression according to the experimental workflow shown in Fig. 1a.

We first compared the evoked responses obtained with the two spike detection methods. Here, input patterns were encoded by LC, which provided higher classification accuracy (see previous section). The raster plots showed that the temporal arrangement of the stimulus-locked responses was largely similar between the two methods, with response patterns recurring at intervals of approximately 100 ms in both cases (Fig. 4a). Quantitative analysis revealed that the mean firing rate during stimulation was significantly lower with spike sorting, and the number of units was also significantly reduced (Fig. 4b). In contrast, the effective rank of the evoked responses did not differ significantly between the two methods. These results suggest that spike sorting removed redundant spikes recorded by the HD-MEA and reduced the number of active units while largely preserving the dimensional structure of population activity.

**Fig. 4.**
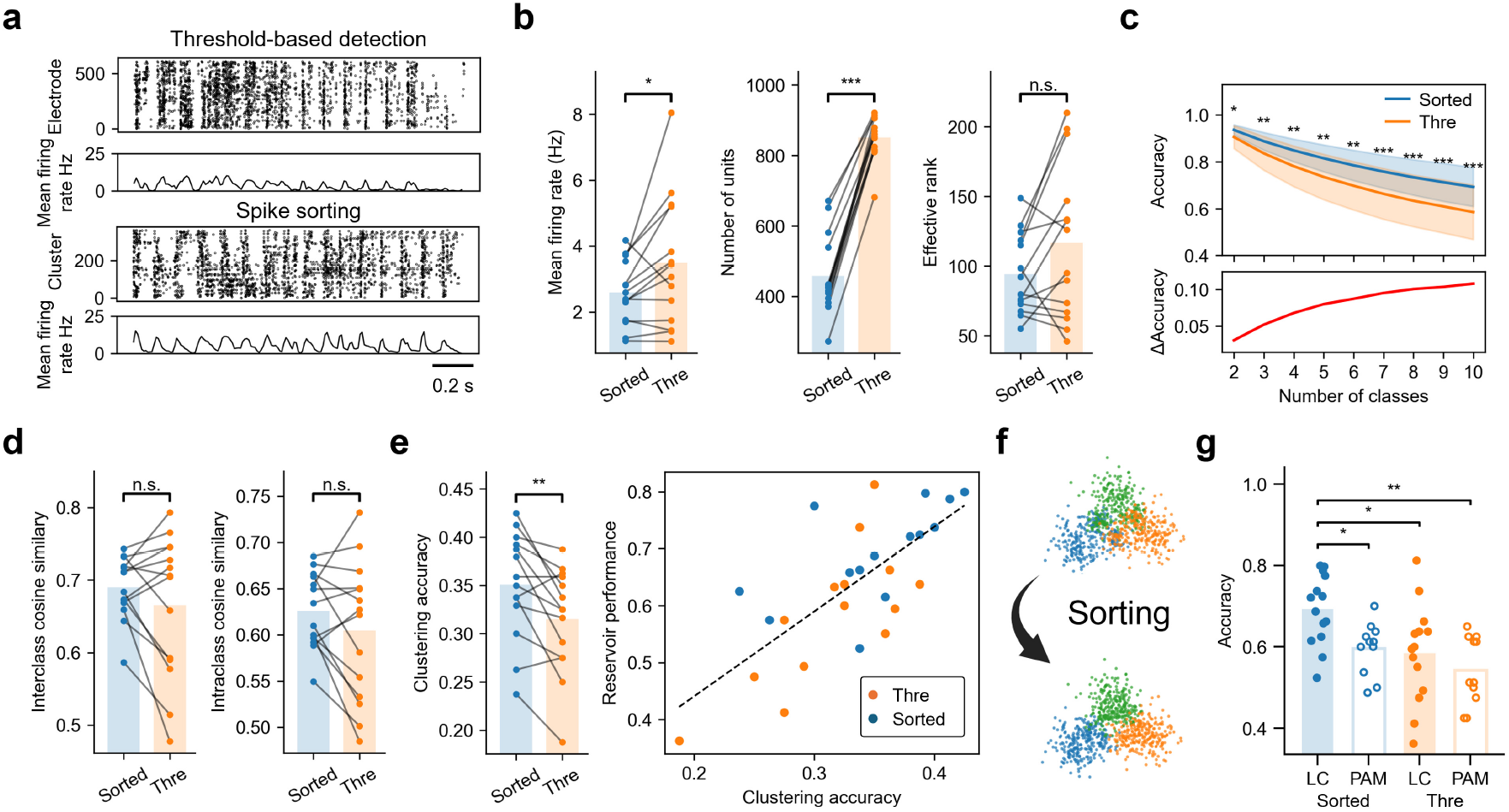
Effects of detection methods on reservoir computing. (a) Representative spike rasters and firing-rate traces during stimulus-evoked responses obtained from threshold-based detection (Thre; top) and spike sorting (Sorted; bottom). Latency coding was used for input encoding. (b) Comparison of firing rate (left), number of units (middle), and effective rank (right) between two detection methods. Number of units corresponds to the number of recording electrodes for threshold-based detection and clusters for spike sorting. For each metric, bars represent the mean across samples. \**p <* 0.05; \*\*\**p <* 0.001; n.s., not significant (two-sided paired *t* test). (c) Classification accuracy as a function of the number of input classes. Orange and blue lines indicate threshold-based detection and spike sorting, respectively (top). The difference in mean accuracy between spike sorting and threshold-based detection is shown in the bottom panel. \**p <* 0.05, \*\**p <* 0.01 and \*\*\**p <* 0.001 (two-sided paired *t* test). (d) Intra- and interclass cosine similarities obtained from threshold-based detection and spike sorting. Intraclass similarity was defined as the mean cosine similarity between activity vectors and their corresponding centroid, whereas interclass similarity was defined as the mean pairwise cosine similarity between different class centroids. n.s., not significant (two-sided paired *t* test). (e) The clustering accuracy obtained by two detection methods. Reservoir states were clustered using *k*-means (*k* = 10), and the resulting cluster labels were optimally matched to the true class labels. The clustering accuracy was defined as the accuracy after optimal label matching (left). Relationship between the clustering accuracy and reservoir classification performance. Orange and blue dots indicate threshold-based detection and spike sorting, respectively (right). Dashed lines indicate linear fits for both methods. \**p <* 0.01 (two-sided paired *t* test). (f) Schematic illustration of class separation in the reservoir state space by threshold-based detection (top) and spike sorting (bottom). (g) Comparison of classification accuracy across four combinations of input encoding and spike detection methods in a 10-class classification task. Blue and orange indicate data obtained from spike sorting and threshold-based detection, respectively. Filled and open bars/dots indicate data from LC and PAM, respectively. Bars represent mean values, and dots indicate individual recordings. Only significant comparisons are shown. \**p <* 0.05; \*\**p <* 0.01 (two-sided Welch’s *t* tests with Holm correction for six pairwise comparisons). Sample size: *n* = 14 recordings from 12 networks (LC) and *n* = 11 recordings from 10 networks (PAM) for each detection method.

Evaluation of the classification performance revealed that spike sorting yielded significantly higher accuracy than threshold-based detection did across all numbers of classes (Fig. 4c). Under the 10-class condition, classification accuracy increased from 58.4 ±11.7% to 69.2 ±8.3% with spike sorting. The advantage of spike sorting was smaller for tasks involving fewer classes and increased with the number of classes. Thus, classification performance improved with spike sorting despite the lower mean firing rate and smaller number of units.

To identify the basis of this improvement, we next characterized how the geometry of the activity vectors changed by spike sorting. Comparison of cosine similarities in interclass and intraclass responses revealed that the values did not differ significantly between the two methods (Fig. 4d; *p >* 0.05, two-sided paired *t* test). Instead, the improved classification performance was associated with increased class separability. A *k*-means clustering analysis revealed that the clustering accuracy was significantly higher with spike sorting and was positively correlated with classification accuracy (Fig. 4e; Pearson’s *r* = 0.69, pooled across both detection methods). These results suggest that spike sorting may improve multiclass classification by reducing redundant activity components, thereby enhancing class separability without altering the overall geometry of the activity representations (Fig. 4f).

Finally, we examined how the combination of input encoding and spike detection methods affected the classification performance (Fig. 4g). For LC, the mean accuracy in the 10-class classification task was significantly higher with spike sorting than with threshold-based detection. However, this improvement was not statistically significant for PAM. This could be caused by the difference in the information representation between PAM and LC. In PAM, the evoked neural responses exhibit rate-coding-like properties because input signals are encoded by modulating the stimulation amplitude, which alters instantaneous firing rates across the network. Thus, population-level activity, which can be captured independently of the detection method (Fig. 2d), likely plays an important role in representing input information in PAM. Spike sorting may therefore add little information beyond what threshold-based detection already conveys.

## 4 Discussion

We investigated how information coding strategies influence the performance of BNN-based RC. By systematically comparing input encoding and spike detection methods in a spoken digit classification task, we found that LC combined with spike sorting yielded the highest classification accuracy (Fig. 4g). Importantly, the advantage of these methods was smaller for tasks involving fewer classes and became more pronounced as the number of classes increased.

The relationship between effective rank and classification accuracy differed between the two input encoding schemes. The basic idea of RC is to map input signals into high-dimensional space, thereby enabling even a linear, memoryless decoder to generate appropriate outputs [40]. In contrast, LC-based stimulation yielded higher classification accuracy than PAM-based stimulation, although it produced reservoir states with lower effective ranks (Figs. 3c,e). Because LC embeds input information in the timing of stimulation pulses, these results suggest that reproducible stimulus-locked temporal responses, rather than dimensionality alone, support superior performance in LC. Although the effective rank under LC was lower than that under PAM (Fig. 3e), it was not apparently limiting for the present 10-class classification task. Thus, within the dimensional range examined here, performance may have relied less on dimensionality and more on how effectively neuronal populations encoded input stimuli into reproducible and class-discriminative activity patterns.

The influence of spike sorting on the analysis of neural dynamics and the performance of downstream decoders remains debated. For example, an in vivo study has shown that population-level neural dynamics can be largely preserved even when neural activity is detected using threshold-based detection [41]. In contrast, in vivo neural decoding studies have suggested that precise identification of spike events can improve decoding performance [42]. Our results are consistent with both perspectives, in that the benefit of spike sorting depended on task complexity. Specifically, although the advantage of spike sorting was significant across all numbers of classes, the gain was smaller in simpler tasks with fewer classes. Given that population-level neural activity was largely preserved between threshold-based detection and spike sorting (Figs. 2c,d), these results suggest that threshold-based detection may be sufficient for simple tasks that do not require a fine separability of activity patterns.

We also showed that BNN-based RC achieved a classification accuracy of 69.2% in a 10-class spoken digit classification task. The 10-class temporal classification task is a more demanding benchmark than those used in many previous BNN-based RC studies [18, 19, 21]. Although the tasks differ, the accuracy achieved here exceeds the performance reported in cortical organoid-based RCs before unsupervised improvement through repeated trials [19]. This high performance is attributed not only to the appropriate choice of input encoding and spike detection methods but also to the microfluidic control of the neuronal network structure [32]. Introducing modular structures into BNNs suppresses globally synchronized activity and diversifies neural dynamics [31, 38], a property that has been shown to support temporal pattern generation in BNN-based RCs [20]. Although the present study did not directly compare patterned and nonpatterned networks, such microfluidic structural control may also have contributed to the high classification performance observed here.

As MEA technology continues to advance through higher electrode density and greater flexibility in stimulation control, BNNs will become increasingly accessible as precisely controllable computational substrates. In this context, input encoding schemes should be regarded not only as methods for delivering stimuli but also as key design parameters for shaping reservoir states. The present study provides practical guidelines for designing future BNN-based RC systems. Future studies should further clarify how input encoding and spike detection methods interact with long-term plasticity induced by repeated stimulation [19, 21] and how these coding strategies operate in closed-loop settings that incorporate sensory feedback from dynamical environments [43]. Real-time implementations of spike sorting enabled by hardware accelerators will also be important for closed-loop RC applications [44]. Such knowledge is essential for harnessing the full potential of BNNs as physical reservoirs.

## Acknowledgements

We also thank Mr. Iori Morita at Tohoku University for the photomasks. This work was partly supported by MEXT Grant-in-Aid for Transformative Research Areas (A) “Multicellular Neurobiocomputing” (24H02330, 24H02332), JSPS KAKENHI (23H02805, 23K28179, 25K24740, 26H02512, 26K21942), JST CREST (JPMJCR19K3), JST ALCA-Next (JPMJAN23F3), the WISE Program for AI Electronics of Tohoku University, and the Cooperative Research Project Program of the RIEC, Tohoku University. This research was carried out at the Laboratory for Nanoelectronics and Spintronics, RIEC, Tohoku University.

